# Physical model of electrical synapses in a neural network

**DOI:** 10.1101/2021.12.13.472453

**Authors:** M. Pekker, M.N. Shneider

## Abstract

A theoretical model of electrical synapses is proposed, in which connexons play the role of “nails” that hold unmyelinated areas of neurons at a distance of about 3.5 nm, and the electrical connection between them is provided by charging the membrane of an inactive neuron with currents generated in the intercellular electrolyte (saline) by the action potential in the active neuron. This mechanism is similar to the salutatory conduction of the action potential between the nodes of Ranvier in myelinated axons and the ephaptic coupling of sufficiently close spaced neurons.

## I. Introduction

An electrical synapse (Fig. 1) is similar to the connection of wires in an electrical circuit, which allows, in contrast to a chemical synapse, the propagation of an action potential in both directions, that is, both from neuron A to neuron B and in the opposite direction [1–6]. In works [2–6], it is assumed that the electrical connection between neurons is carried out due to the transport of ions and small molecules through the connexons. Various options for connecting neurons with electrical synapses (more precisely, by connexons) considered in the literature are shown in Fig. 1. The mathematical description of electrical synapses is reduced to an electrical circuit connecting two axons, along one of which action potential propagates [7–12]. As a rule, the simplified equivalent electrical circuit does not reveal the physical mechanism underlying the operation of the electrical synapse and is phenomenological in nature.

**Figure 1.**
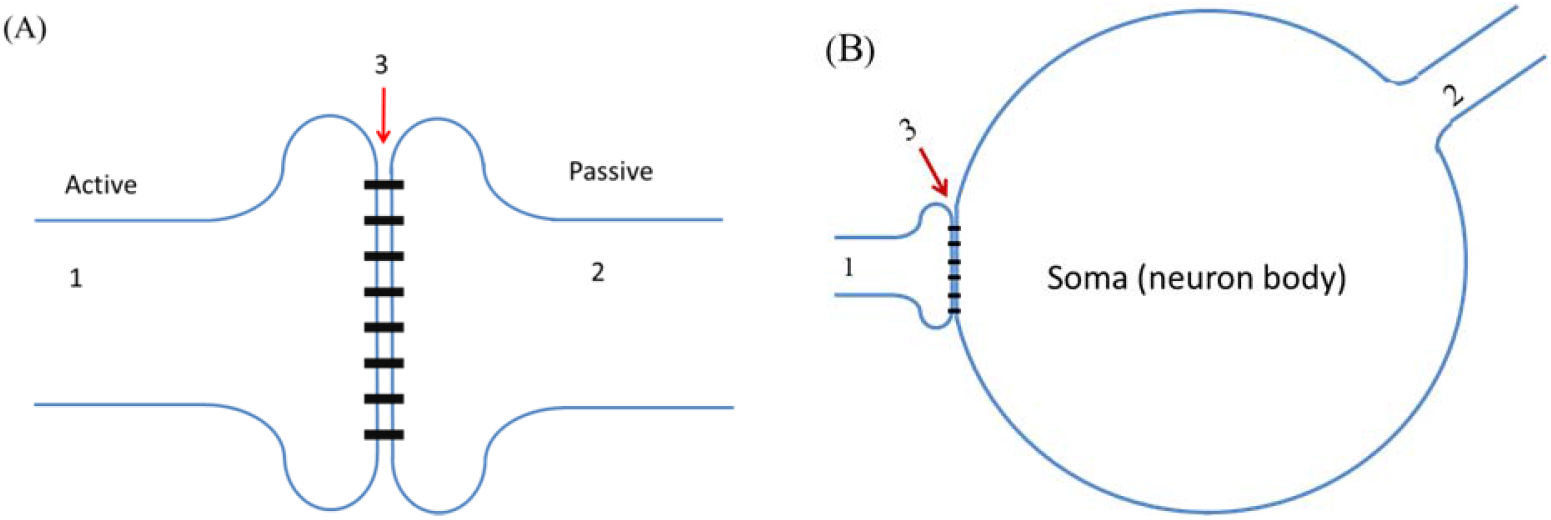
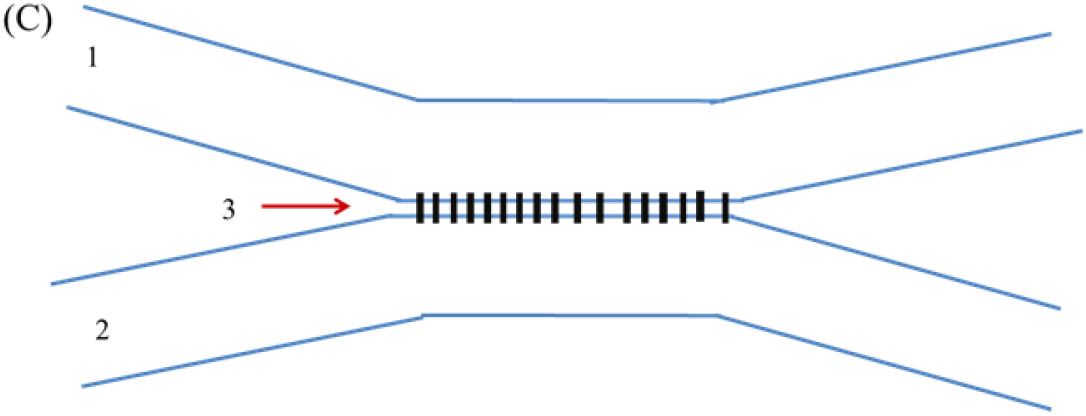
Electrical synapse. A – Connecting an axon to a dendrite (or other axon) through an electrical synapse. 1 – Active axon, 2 – Dendrite (or passive axon), 3 – A system of connexons. The action potential propagating along axon 1 “jumps” through the electrical synapse to the dendrite (or axon 2) and then spreads in it. B – Electrical connection by connexons of the axon with the soma of the neighboring neuron. An action potential propagating along axon 1, through an electrical synapse, excites an action potential on the body of a neighboring neuron, which in turn stimulates the propagation of an action potential in axon 2. C – Connection of non-myelinated regions of two axons 1 and 2 by connexons 3.

In this work, we consider a model of electrical synapse with connexons that hold non-myelined areas of neurons at a distance of about 3.5 nm (see, for example, [13, 14]). We will show that the transfer of excitation between neurons occurs due to the charging of the membrane of an inactive neuron with currents generated in the intercellular saline by the action potential in the active neuron. Furthermore, the mechanism of transmission of excitation in the electrical synapse is essentially similar to the synchronization mechanism between nearby neurons (ephaptic coupling) considered in [15, 16].

## II. The dynamics of potential and currents in the vicinity of a neuron in the case of the propagating action potentials

Saline, the fluid in which the neurons are located in living organisms, is a highly conductive electrolyte, with *σ*~1 – 3 Ω^-1^*m*^-1^ [16, 17]. Therefore, the relaxation time of the volume charge in it (the Maxwell time) is of the order 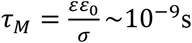. [18], where *ε*_0_ is the dielectric constant of the vacuum, and *ε* ≈ 80 is the relative dielectric permittivity of water, i.e., by 5–6 orders of magnitude shorter than the characteristic time of the excitation and relaxation of the action potential in the axons. The size of the non-quasi-neutral region near the membrane of axons is determined by the Debye length 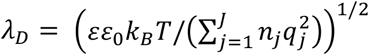. Here *k_B_* is the Boltzmann constant, *T* is the temperature, and *n_j_* is the densities of the ions with the charges *q_j_*. In the case of saline, we can assume that all the ions in the liquid are singly charged and that their total density is of the order *n_j_* ≈ 2·10^26^m^-3^ [19]. For *T* ≈ 300 *K*, *λ_D_* ≈ 0.5 *nm*, and this value is much smaller than the typical radius of the axon *R*_0_ ≈ 1 – 8 μm. Therefore, the violation of quasi-neutrality cannot be taken into account, and the potential distribution in the vicinity of a neuron can be found from the equation of the continuity of current

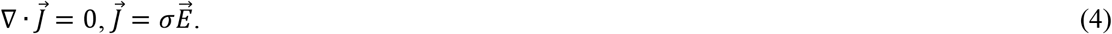

Since the conductivity *σ*of the electrolyte is constant, and 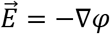, the problem of determining the potential is reduced to the Laplace equation:

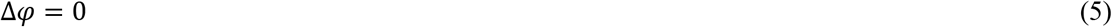

For definiteness, consider two cases of electrical synapse. The first case is when an electrical synapse connects the axon to the dendrite (Fig. 1A) and the second to the neuron body (soma) (Fig. 1B). Since the distance between the soma and the axon is about *d_s_* =3.5 nm, the radius of the axon is *R*_0_~1-8 μm [20], and the radius of the neuron body (soma) *R_b_* is several times larger than the radius of the non-myelinated region of the axon, then *R_b_* ≫ *R*_0_ ≫ *d_s_*. In the case of the electrical contact of an axon with a dendrite or soma, the axon can be considered a cylinder, with a radius of *R*_0_, directly connected to a cylindrical dendrite (axon) or to the soma of a neighboring neuron. Since *R_b_* ≫ *R*_0_, then, without loss of generality, the body of the axon can be considered a plane.

The estimation of the charging time of a cylindrical capacitor of radius *R*_0_ and capacitance per unit area *c_m_* in an electrolyte with conductivity *σ* is [21]:

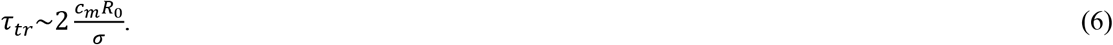

Since the membrane capacity of the unmyelinated section of the axon per unit area is 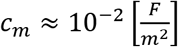, and the electrolyte conductivity *σ* ~ 1-3 ⋂^-1^m^-1^ [17], then for an axon radius of 1.5-8 μm [21], *τ_tr_* varies from 0.035 to 0.1 μs. According to [17], the typical duration of the action potential impulse at the interface between the active axon and the passive dendrite (passive axon) is *τ_pa_*~1 ms. Since *τ_tr_* ≪ *τ_pa_*, we can assume that each value of the radial current on the active axon corresponds to the acquisition of an additional potential on the membrane of the passive axon. The same applies to the contact of the axon with the soma since the radius of the neuron is only several times the radius of the axon [17]. Obviously, the statement about the linear connection of the potential on the inactive axon or on the body of the neuron from the radial current on the active axon is valid until the additional potential on the dendrite or soma of the neighboring neuron exceeds the threshold value of the initiation of the action potential. In other words, up to this point in time, the membrane of a dendrite, axon, or soma of a neighboring inactive neuron can be considered impermeable to ions. Accordingly, the boundary condition on an inactive dendrite (axon) (Fig. 2A) is:

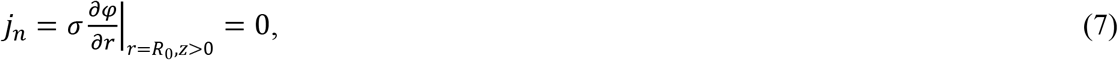

and on the soma of the neuron (Fig. 2B):

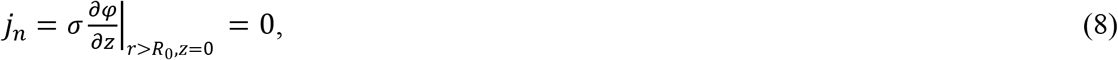

where *j_n_* is the normal component of the current on the dendrite (axon) or soma of an inactive neuron.

**Figure 2.**
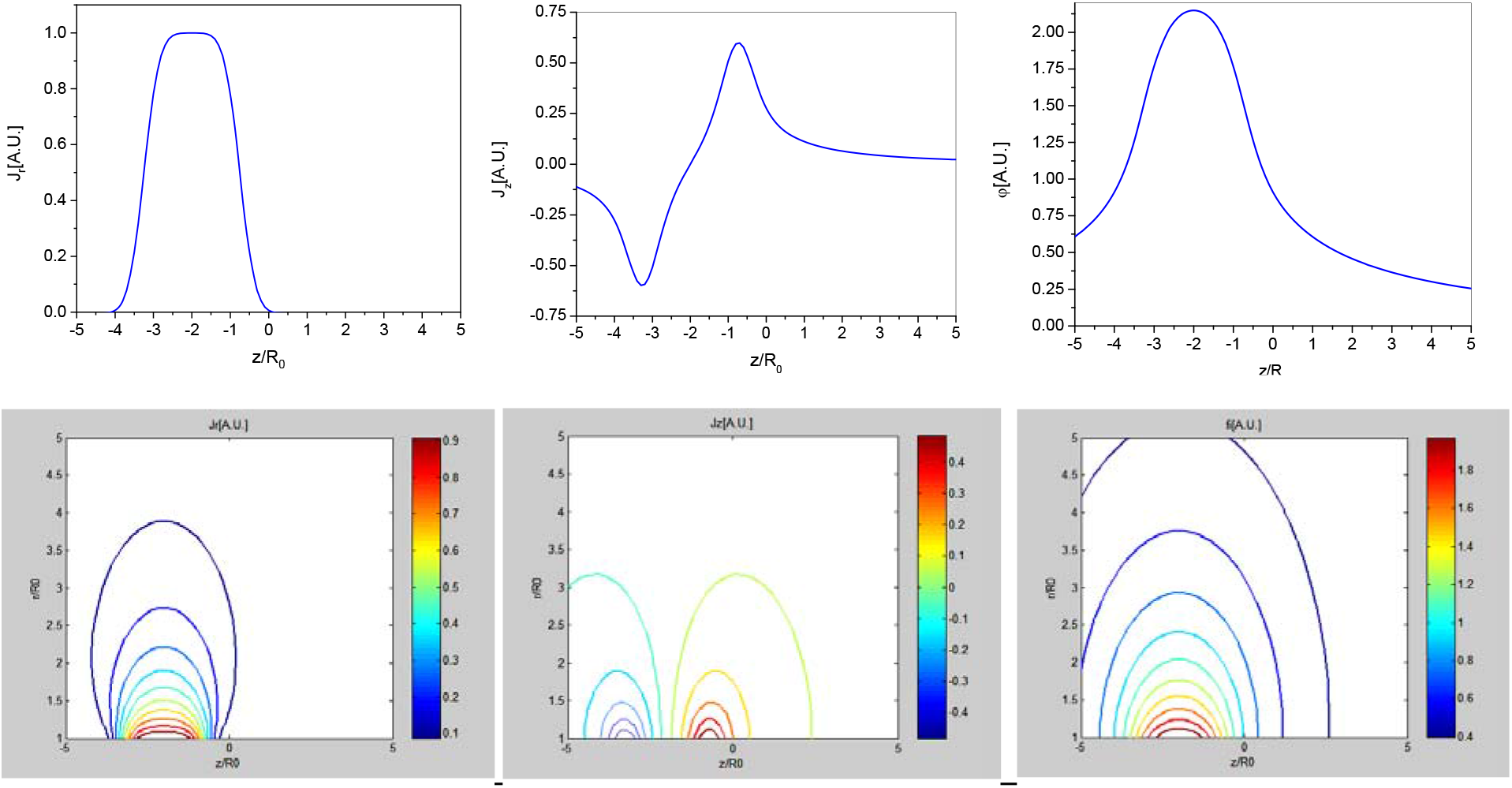
Current and potential distributions for a rectangular radial current pulse on contiguous axons. (A), (B), (C) – Distributions of radial current 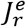, longitudinal 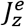 and potential *φ^e^* on the membranes of contiguous neurons, (D), (E), (F) – Corresponding volumetric distributions of currents and potential in saline. The active axon corresponds to the line *r*/*R*_0_ = 1, *z* < 0, the passive one – *r*/*R*_0_ = 1, *z* > 0. Presented calculation results correspond to *b* = 2/*R*_0_, *z_i_*/*R*_0_ = –1, –1.5, –2, –2.5, –3, *J*_1_ = *J*_2_ = *J*_3_ = *J*_4_. The currents 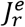 and 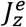 are normalized to the amplitude of the radial current *J_am_* on the surface of the active neuron and the potential *φ^e^* to *φ_norm_* = *J_am_R*_0_/*σ_e_*.

If we assume that near the electrical contact, the radial current through the membrane has a Gaussian spatial distribution and does not depend on time,

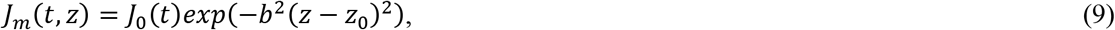

where *J*_0_(*t*) is the maximal current through the membrane, then the distribution of the potential *φ^e^* and currents outside the axon have the form [15, 16]:

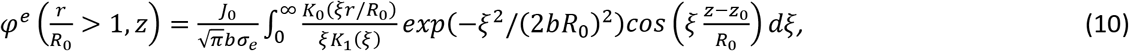

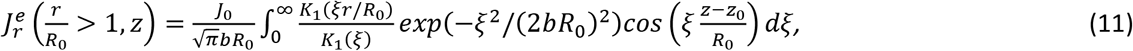

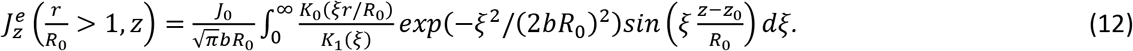

We can say that formulas (10)-(12) determine the corresponding distributions of the potential and currents in the vicinity of the contacts of electrical synapses shown in Fig. 1.

It should be noted that in works [22–24], a different approach was used to calculate the potential and currents outside the axon. In these works, it was assumed that not the current through the membrane of the active axon was given but the potential difference on the inner and outer sides of the membrane.

It is obvious that the electrical contact of an active axon with the soma of a neighboring unexcited neuron can be described by a radial current:

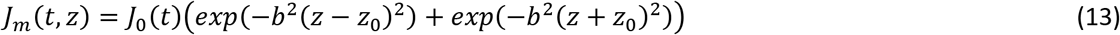

The distribution of the potential and currents corresponding to the current (13) in the adjacent regions of the intercellular electrolyte has the form:

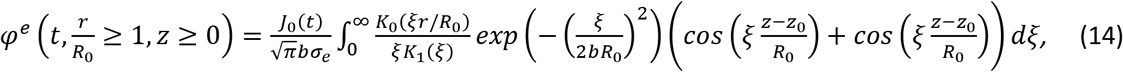

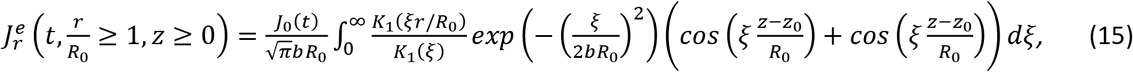

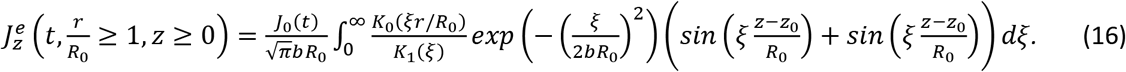

In general, any distribution of radial currents on the membrane of the active axon can be approximated by the sum of Gaussian distributions:

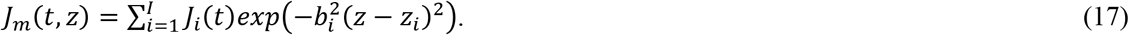

In this case, when (17) is substituted into (10) - (12), it is easy to obtain formulas for describing the electrical contact of an active axon with a dendrite and when (17) is substituted into (14) - (16) for the electrical contact of an axon with the soma of a neighboring inactive neuron.

Since the values of the currents and potential on the passive dendrite (or soma) are proportional to the radial current on the membrane of the active neuron *J_m_*, in the calculations we assumed *J_m_* to be time independent.

Fig. 2 shows the distributions of the currents and potentials on the membrane of active and passive axons and in the electrolyte region for the case of contact активного аксона с пассивным дендритом или аксоном.

Fig. 3 shows the distributions of the currents and potential on the soma membrane of an unexcited neuron and in the adjacent areas of the electrolyte.

**Figure 3.**
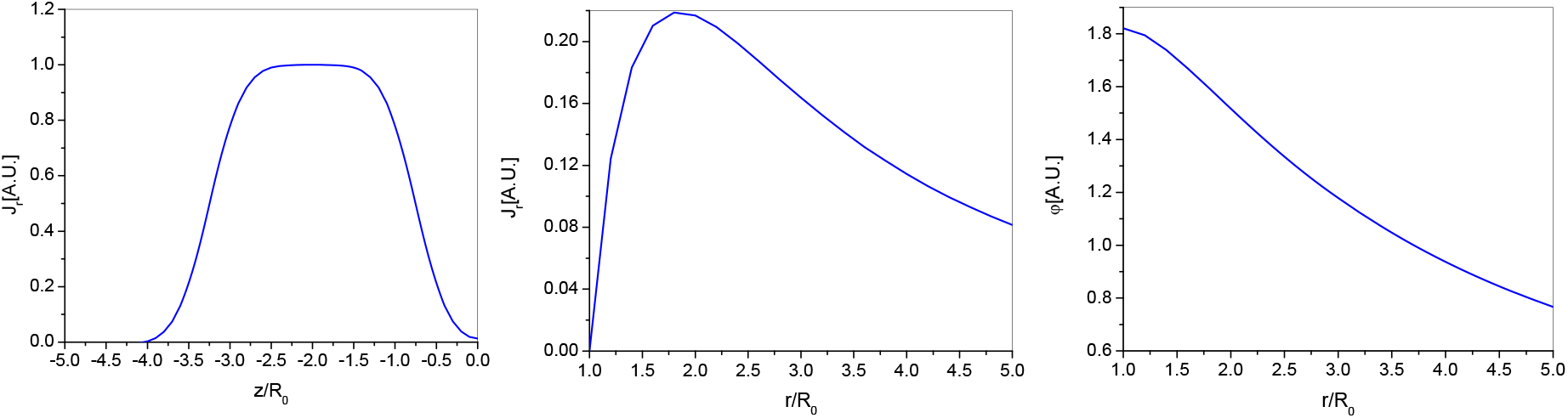
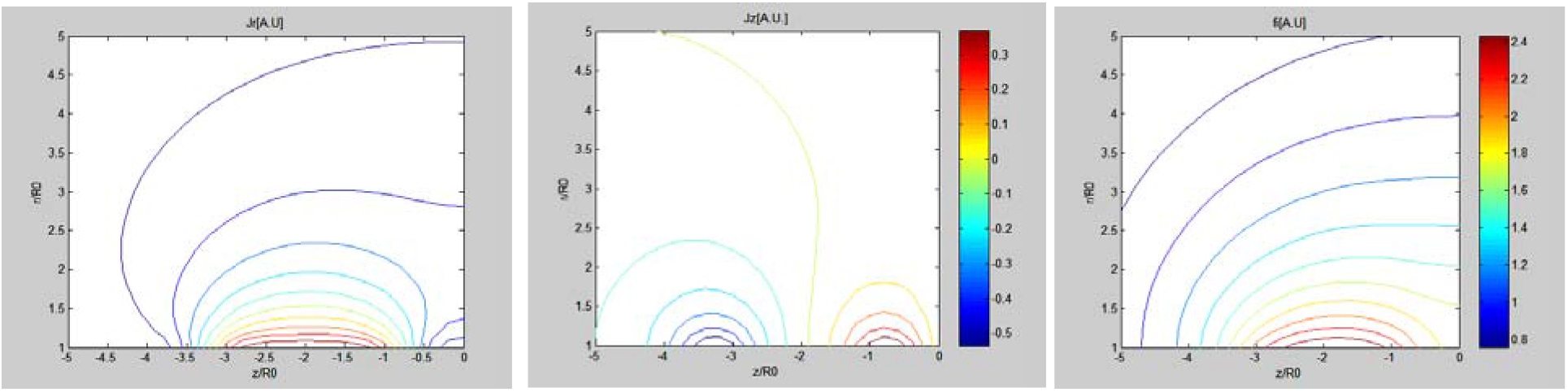
Current and potential distributions for a rectangular radial current pulse on an active axon in contact with soma (see Figs. 1B and 2B). (A) – Distribution of radial current 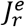 on the membrane of the active axon, (B) and (C) – Distributions of radial current 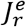 and potential *φ^e^* on the membrane of the soma of a passive neuron; the longitudinal current on soma равен нулю. (D), (E), (F) – Corresponding volumetric distributions of currents and potential in saline. The line 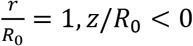 corresponds to the active axon, the line 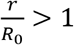, and *z*/*R*_0_ = 0 crresponds to the soma. Presented calculation results correspond to *b* = 2/*R*_0_, *z_i_*/*R*_0_ = –1, –1.5, –2, –2.5, –3; *J*_1_ = *J*_2_ = *J*_3_ = *J*_4_. The currents 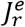 and 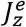 are normalized to the amplitude of the radial current *J_am_* on the surface of the active neuron and the potential *φ^e^* to *φ_norm_* = *J_am_R*_0_/*σ_e_*.

As follows from the calculations, the characteristic size of the change in the currents and potential on the passive axon and the neuron body is of the order of 2–3 axon radii. Therefore, neglecting the size of the gap between the active axon and the dendrite or soma and considering the soma as a plane is acceptable in our calculations. At the same time, calculations show that the change in potential on the surface of the dendrite (axon) or the surface of the soma of an unexcited neuron is up to 30% of the maximum potential on the active axon. That is, the potential difference at the membrane surface (external and internal) in a passive dendrite (axon) or soma can be up to 20–30 mV. In principle, this should be sufficient to activate the action potential via the electrical synapse.

Note that the considered mechanism of the “jumping” of the action potential from the active axon to the passive one through the electric synapse gap is also applicable at the surface contact shown in Fig. 2C. Thus, the role of connexons is reduced to keeping axons and dendrites close enough for the action potential to “jump” from the active site to the passive one.

## Conclusions

A model of electrical synapse has been proposed in which connexons play the role of “nails,” holding non-myelined areas of neurons at a distance of several nanometers. In this case, the transmission of excitation in the electrical synapse is provided by charging the membrane of an inactive neuron with currents generated by the action potential in the saline by an active neuron. This model explains the transfer of excitation across the electrical synapse without the need for generating ion fluxes across connexons.

